# A head-fixation system for continuous monitoring of force generated during behavior

**DOI:** 10.1101/868703

**Authors:** Ryan N. Hughes, Konstantin I. Bakhurin, Joseph W. Barter, Jinyong Zhang, Henry H. Yin

**Affiliations:** Department of Psychology and Neuroscience, Duke University, Durham, NC, 27708, USA; Department of Neurobiology, Duke University School of Medicine, Durham, NC, 27708, USA

## Abstract

Many studies in neuroscience use head-fixed behavioral preparations, which confer a number of advantages, including the ability to limit the behavioral repertoire and use techniques for large-scale monitoring of neural activity. But traditional studies using this approach use extremely limited behavioral measures, in part because it is difficult to detect the subtle movements and postural adjustments that animals naturally exhibit during head fixation. Here we report a new head-fixed setup with analog load cells capable of precisely monitoring the continuous forces exerted by mice. The load cells reveal the dynamic nature of movements generated not only around the time of task-relevant events, such as presentation of stimuli and rewards, but also during periods in between these events, when there is no apparent overt behavior. It generates a new and rich set of behavioral measures that have been neglected in previous experiments. We detail the construction of the system, which can be 3D-printed and assembled at low cost, show behavioral results collected from head-fixed mice, and demonstrate that neural activity can be highly correlated with the subtle, whole-body movements continuously produced during head restraint.

## Introduction

Many studies in modern behavioral neuroscience use head-fixation (Toda et al., 2017;Coddington and Dudman, 2018;Economo et al., 2018). Historically, head-fixed experimental preparations in awake behaving animals have contributed to the study of motor control (Evarts, 1968;Hikosaka and Wurtz, 1983), associative learning (Waelti et al., 2001;Eshel et al., 2015), and even navigation (Dombeck et al., 2010). Head-fixation confers two major advantages. First, it allows convenient monitoring of neural activity that minimizes noise and motion artifact. This enables the use of powerful techniques for recording neural activity on a large scale, such as two-photon calcium imaging or electrophysiology using large electrode arrays. Second, by greatly constraining the behaviors that the animal can perform, it simplifies experimental design and data analysis. For this reason, studies using head-fixation usually focus on the behavior of interest, and the typical measures include licking, EMG measures of arm or mouth muscle activity, bar pressing, and eye movements (Newsome et al., 1989;Schultz et al., 1992;Eshel et al., 2015;Economo et al., 2018). While these measures have provided valuable insights, other behaviors generated by the head-fixed animals are neglected. For example, many postural adjustments of the body are not measured. Researchers often assume that no other behaviors, such as head movements, are occurring when animals are head-fixed. However, just because these subtle movements cannot be easily observed, it does not follow that there are no attempts to move the head or other body movements. Furthermore, often animals are placed in chambers or rooms away from the experimenter, so that close observation of the actual behavior is not even attempted. Because neural activity recorded during standard behavioral tasks could well be related to the generation of such movements, they produce significant confounding variables. The lack of knowledge of such variables can therefore result in misinterpretation of neural activity while the true relationships between neural activity and behavior is overlooked.

More recent systems have attempted to address some of these concerns by incorporating a running ball for continuous locomotion in head-fixed animals (Dombeck et al., 2010;Engelhard et al., 2019). During such experiments, animals often make direction-specific adjustments of the body and the head, in addition to their limb movements, in order to steer the ball appropriately. However, the coordinate (x, y, and z) positions of the ball provide little information about the actual kinematics of the body. Others have incorporated accelerometers into their head-fixation devices, thus allowing measurements of whole-body movements (Coddington and Dudman, 2018). The major limitation of accelerometer baskets is they cannot distinguish between movements in different directions, as movements in any direction can result in the same accelerometer reading. Moreover, head movements are not usually measured.

Here we describe a novel head-fixation apparatus equipped with load cells for detection of forces generated by the head and body in head-fixed mice. The system uses multiple sensors to monitor not only the magnitude but also direction of forces that result from head movements as animals move and adjust their posture, revealing the subtle movements that are continuously generated.

## Results

To quantify direction-specific and continuous movements in the head-fixed preparation, we developed a novel head-fixation device that incorporates 5 load cells to directly measure the forces generated by the mice’s head and body movements (Figure 1A, for detailed instructions on assembling the device, see assembly instructions in supplementary materials and exploded diagrams). Mice are implanted with a steel head post as part of routine surgery to implant electrodes into the brain. The post is secured by small pinch clamps, which are part of a lightweight, rectangular plastic frame. The frame is suspended by an array of 3 aluminum load cells, which simultaneously register the forces exerted by the animal in 3 directions (Figures 1A & 1B).

**Fig. 1.**
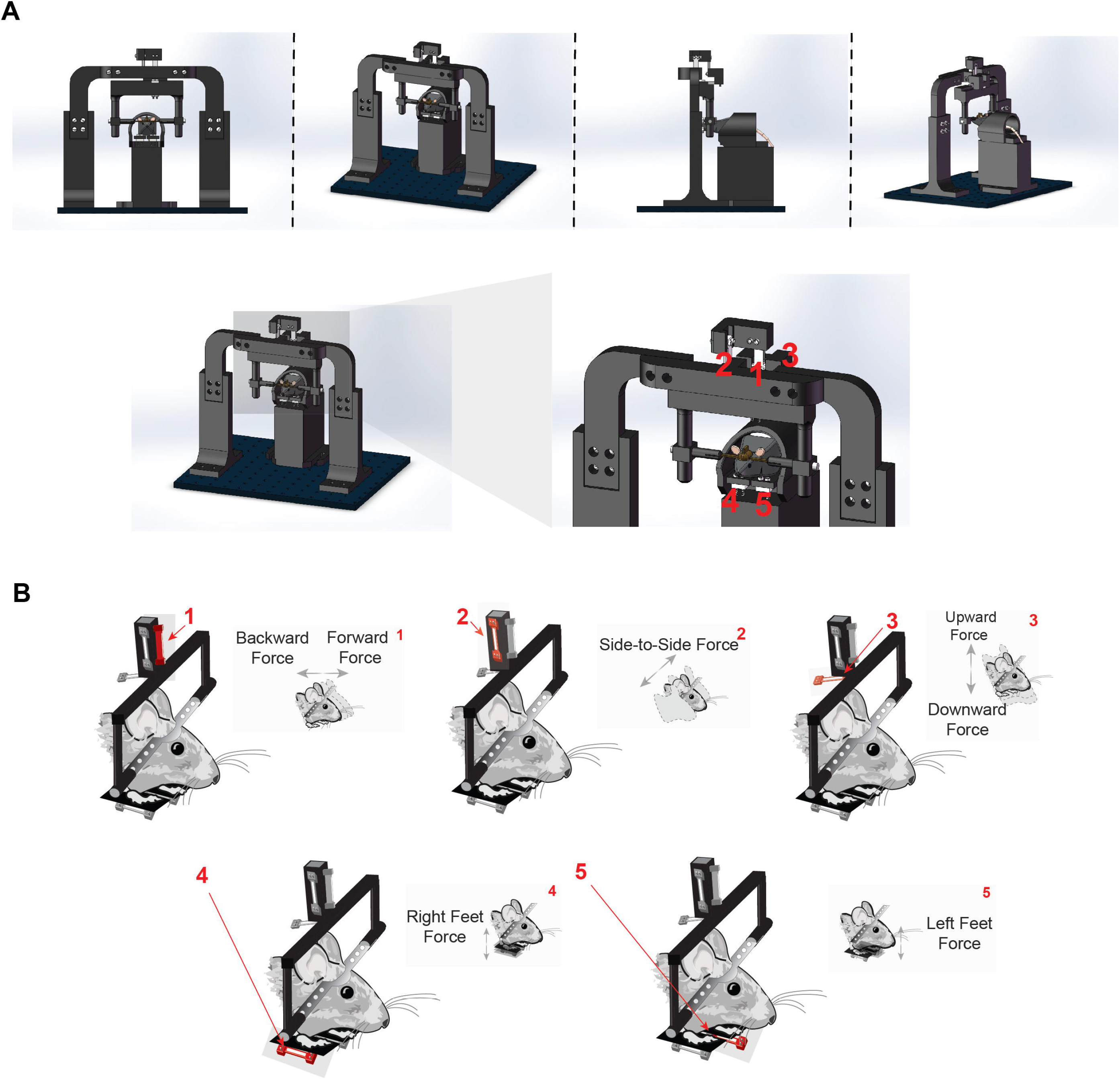
Novel head-fixation set up measures force of the head in 3 orthogonal directions and left and right body forces. **A)** Illustration of novel head-fixation apparatus with five orthogonal force sensors from the front (*left*), side (*middle*) and back (*right*). There are three force sensors that measure the force exerted by the mice’s head (1, 2 and 3). Two additional force sensors are below the mice’s feet on the left and right side of the body (4 and 5). For further instructions, see methods section on assembling the device. **B**) Schematic representation of each force sensors and the direction of force that it measures. Force sensor 1 measures force made by the head in the forwards and backwards direction. Force sensor 2 measures force made by the head in the side-to-side (left and right) direction. Force sensor 3 measures force made by the head in the upward and downward direction. Force sensors 4 and 5 measure force exerted in the upward and downward direction made by the right and left feet, respectively

This precise arrangement of load cells suspending the mouse’s head enabled the measurement of force along orthogonal axes of movement (up/down, left/right, or forward/backward). The load-cells are arranged in a tight cluster together just above the mouse to avoid spurious measures of torque as well as to capture as much of the individual force vector as possible. Two additional sensors were placed below the left and right feet in a custom-designed perch (Figure 1B), in order to measure the forces exerted by each side of the body as the mouse adjusts its posture. All components of the head-fixation device were custom-designed and 3-D printed in the lab.

Small strain gauge load cells designed to operate on a scale of 0-100g were used. Load cells of this type and size are most commonly found in small weighing devices such as kitchen scales. Attached to the aluminum body of the load-cell are sensitive strain gauges arranged in a Wheatstone bridge configuration (Figure 2A). Changes in the shape of the bridge can alter its resistance and voltage levels across the gauges. Because the load-cells register changes to the shape of their aluminum bodies, the cluster of 3 load cells that support the head-fixation frame acts as the frame’s pivot-point as it slightly moves along 3-dimensions in response to the mouse’s movement. The voltage between the Wheatstone bridges is modulated by deformations of the strain gauges, which is proportional to the amount of force exerted. The load cell is coupled to an amplifier circuit that scales this voltage between 0 and 5V, which can be used by most analog recording systems (Figure 2A). The configuration of the circuit shown is designed to detect bidirectional forces exerted on the load cell (Figure 2B). By measuring the slope between known mass and the voltage reading, one can convert the measured voltage to force applied to the load cell. Using a low-resistance trimmer potentiometer between pins 8 and 9 in the amplification circuit, the user can adjust the slope of the voltage change in response to force changes.

**Fig. 2.**
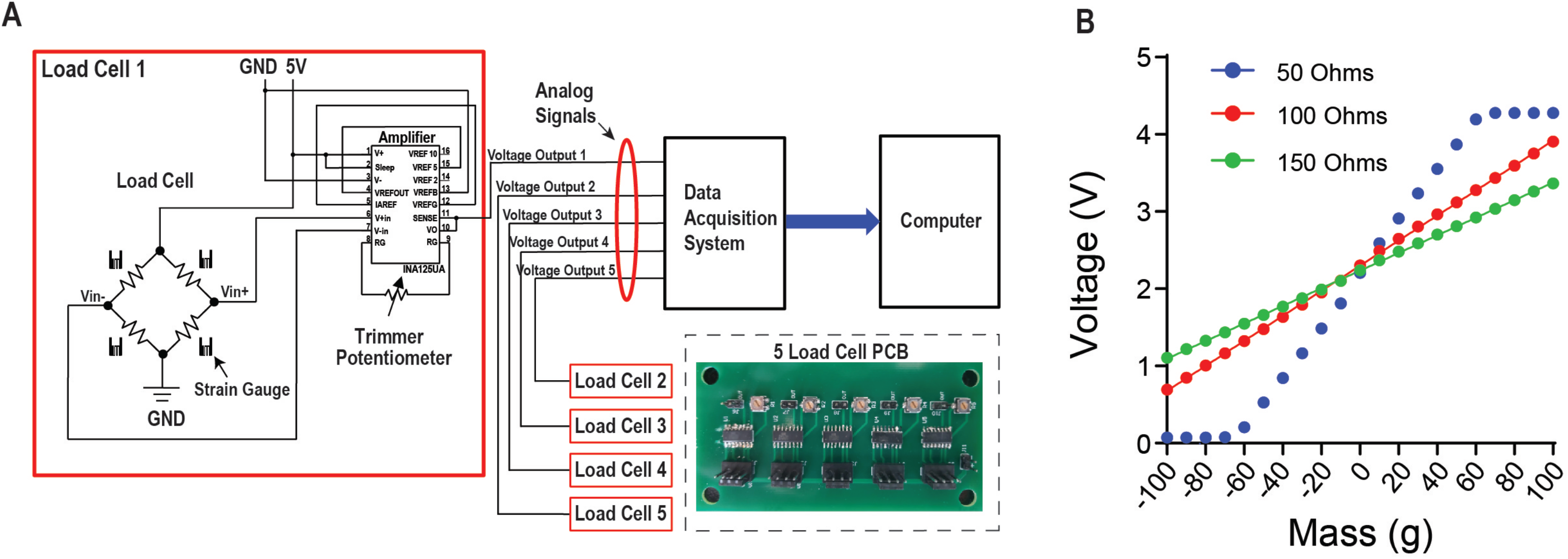
Load cell circuit and set up allows for data acquisition with any device that can read an analog signal. **A)** Circuit diagram for the load cell. The load cell is attached to an amplifier that contains a potentiometer. This allows the user to adjust the slope of the load cell voltage output in response to force changes, and to tune it for the optimal linear relationship between voltage and the force applied. From this circuit, the voltage output is then connected to a data acquisition system to collect continuous voltage signal data. Inset shows the printed circuit board (PCB) designed for 5 load cells. **B**) There is a linear relationship between mass applied to the load cell and the change in voltage output. By using an object with a known mass, a mass-voltage relationship can be obtained using different levels of resistance from the potentiometer.

### Behavior on a fixed-time reinforcement schedule

We tested our load-cell head-fixation device using 2 mice chronically implanted with a head bar. Mice began training on a fixed-time (FT) schedule of reinforcement where they received a 10% sucrose solution every 10 seconds (Figure 3A). During early sessions in the fixed-time task, mice will readily collect water from the spout, but have not yet begun to time the arrival of this drop (Figure 3A). Our system was able to detect the fine and subtle movements during the task. For a given single trial, force exertion was predominantly coincident with their licking behavior after reward delivery, reflecting forward movements towards the spout (Figures 3B & 3G). However, their movements from side to side were different, with one mouse showing large rightward movements and the other showing large leftward movements following the reward (Figures 3C & 3H). Both mice showed a movement upward as they collected the reward (Figures 3D & 3I). By examining the measurements from the force sensors below the feet, we can see that mouse clearly pressed down with both feet (Figures 3E & 3F) to collect the reward, whereas another mouse preferentially applied force with the left side of its body (Figures 3J & 3K). The signals registered by each load cell were distinct for a given trial, reflecting the complexity and amplitude of coordinated movements generated by different parts of the body (Figures 3B-L). In addition, our system could quantify unique movement patterns in each mouse, even when they are engaged in an extremely simple behavioral task.

**Fig. 3.**
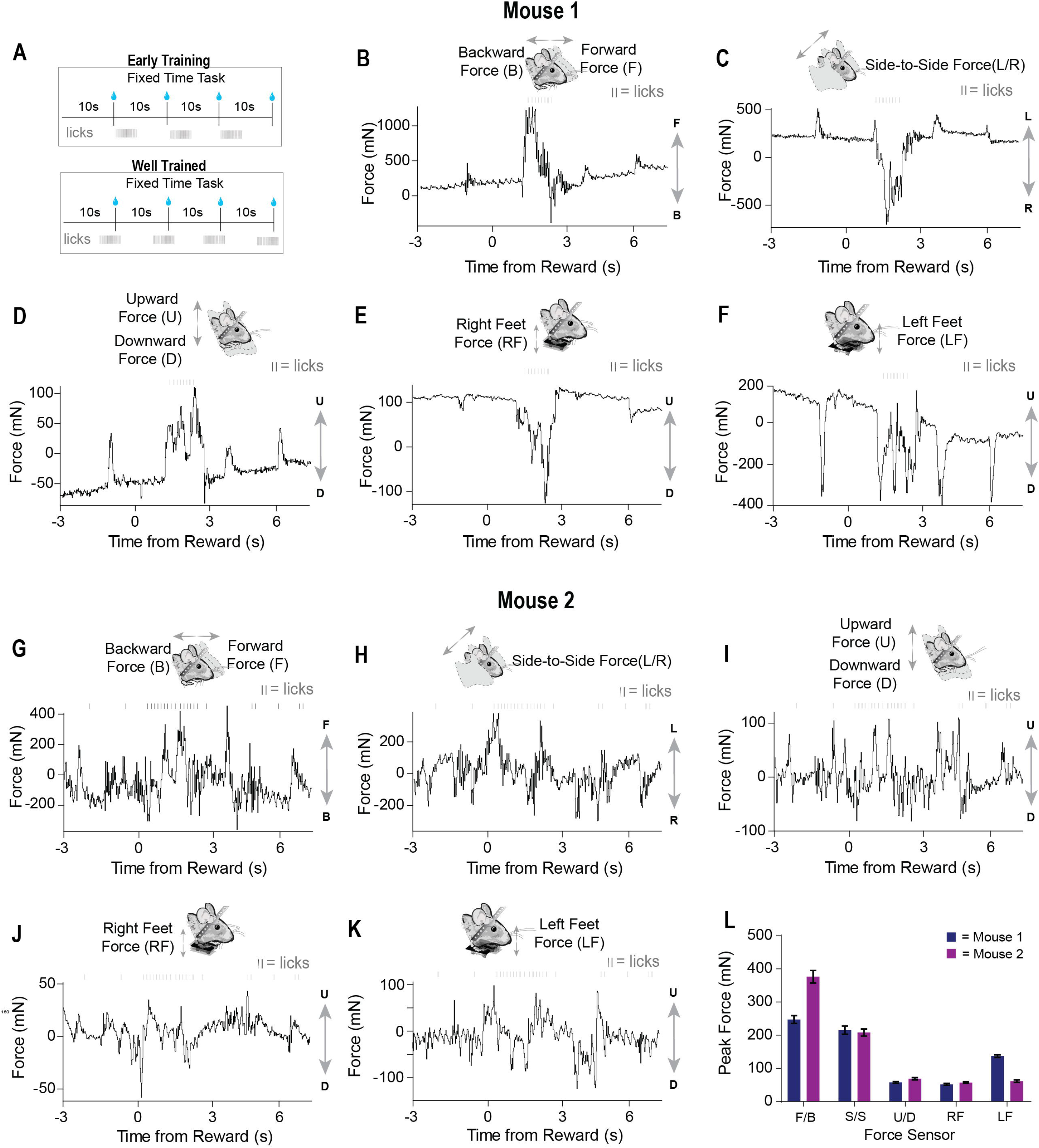
Force sensors detect continuous and fine movements of the head and body during early training in a fixed time (FT) behavioral task. **A)** Two mice were trained on a fixed-time schedule of reinforcement where they received a 10% sucrose solution every 10 seconds. **B-F)** Raw continuous traces of 10 seconds from all five force sensors, as well as lick times, obtained from mouse 1. Each force sensor displays distinct force changes that vary in their amplitude and direction. **G-K)** Raw continuous traces of 10 seconds from all five force sensors, as well as lick times, obtained from mouse 2. **L**) Average peak forces exerted for each force sensor across all trials from both mice.

Well-trained mice (approximately 1-2 weeks) initiated licking before reward presentation (Figure 3A). Similar to what was observed early in training, movements closely coincided with licking. However, after training, both the licking and force measures started before reward delivery (Figure 4). Although both mice displayed stereotyped anticipatory licking before reward (Rossi and Yin, 2015;Rossi et al., 2016;Toda et al., 2017), their movements and postural adjustments were both complex and distinct. Licking-related oscillations were nested within slower movements that reflected postural changes. However, our force measures reveal that, even though the licking becomes stereotyped as the task is learned, the movements from trial-to-trial remain highly variable. In a single reward trial, one mouse displayed forward movements along with large amplitude leftward movements during both anticipatory licking and licking following reward (Figures 4A & 4B). In contrast, the other mouse displayed backward and rightward movements during both anticipatory and consummatory licking (Figures 4G & 4H). Both mice showed upward forces while they licked (Figures 4C & 4I). These upward movements were due to pushing down onto the floor of the perch (Figures 4D, 4E, 4J, & 4K). Although the dynamics of these movements were complex, usually the largest exertion of force by both mice was along the forward/backward and side-to-side axes (Figures 4F & 4L).

**Fig. 4.**
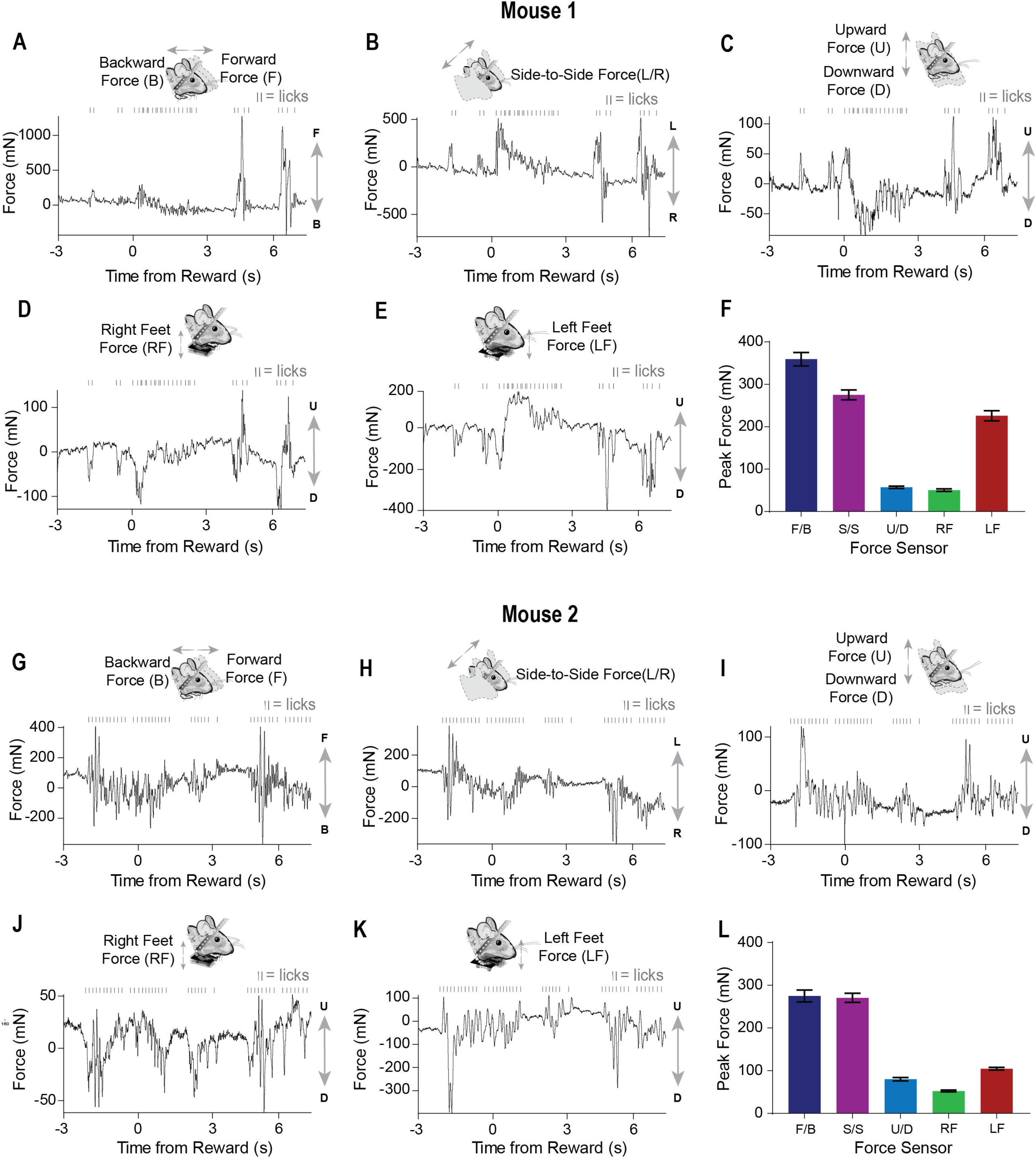
Movements measured by force sensors remain variable even when licking becomes stereotyped when the FT task is well-learned. **A-E)** Raw continuous traces of 10 seconds from all five force sensors, as well as lick times, obtained from mouse 1. Each force sensor displays distinct force changes that vary in their amplitude and direction. **G**) Average peak forces exerted for each force sensor across all trials from mouse 1. **G-K)** Raw continuous traces of 10 seconds from all five force sensors, as well as lick times, obtained from mouse 2. **G**) Average peak forces exerted for each force sensor across all trials from mouse 2.

### Neural recordings of VTA activity while measuring whole body movement forces in head-fixed mice

The dynamic force signals detected by the load-cells reflect subtle changes in head and body posture. To determine the relationship between such continuous measures of movement and neural activity, we recorded single unit activity from the VTA of well-trained mice performing the fixed-time task (*n* = 2, Figure 5A), isolating spiking activity from putative GABAergic neurons in this region. Recent work showed that these neurons can control head angle continuously in freely moving mice (Hughes et al., 2019), but it is unclear how their activity may be related to force exerted by head-fixed mice.

**Fig. 5.**
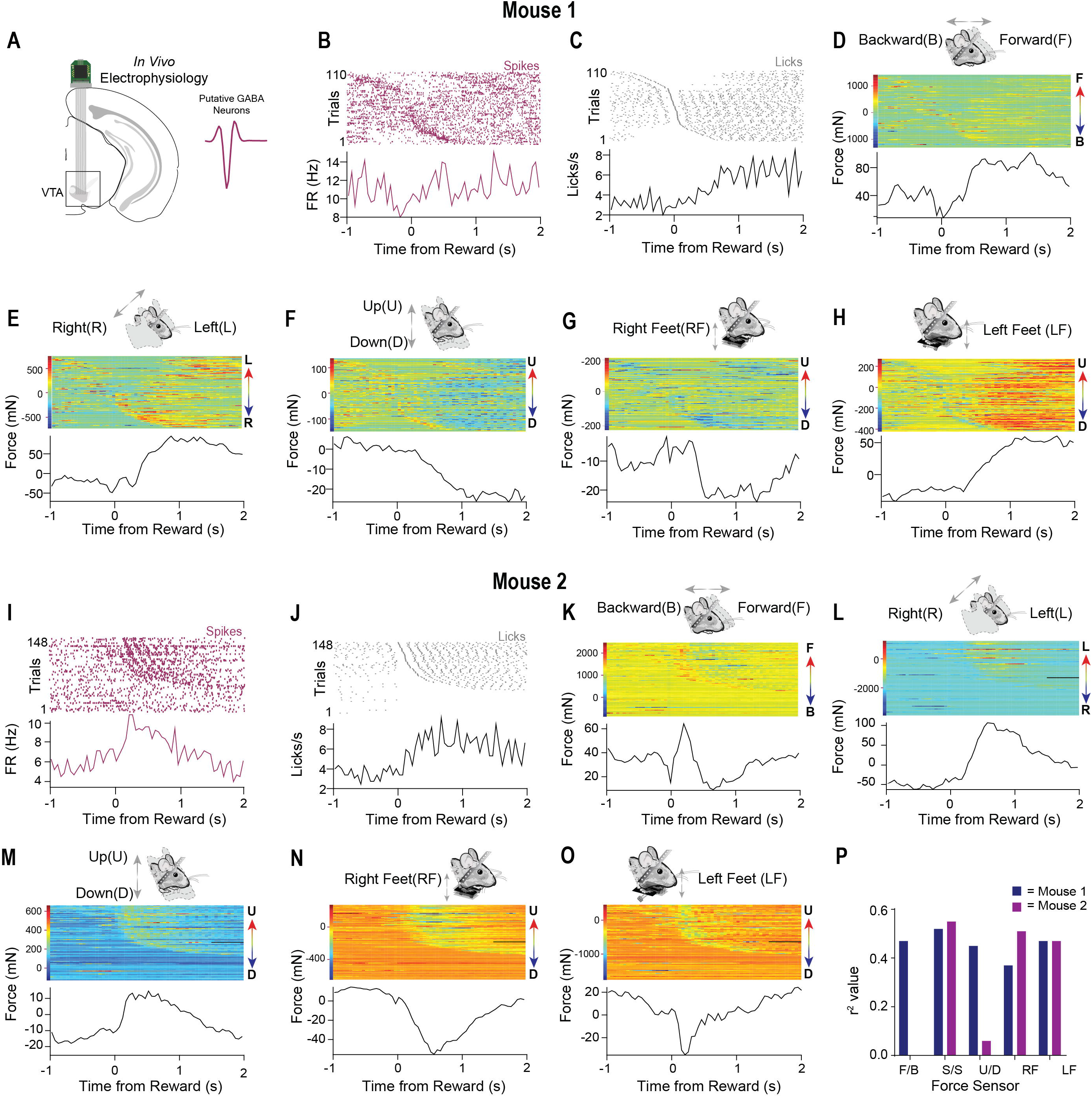
Neural data and precise force measurements can be used to find relationships between neural activity and behavior in head-fixed set ups. **A)** An electrode was implanted into the VTA (n = 2). Waveform of typical putative VTA GABAergic neuron shown on right. **B**) Representative putative GABAergic neuron aligned to the onset of licks after reward from mouse 1. Trials were sorted according to the initiation of the first lick. **C**) Corresponding licks from the same behavioral session. **D-H)** Data from all 5 force sensors aligned to the onset of licks after reward (mouse 1). Color bars on the right of the heatmaps indicate the amplitude and direction of movement in the heatmap (F = forward, B = Backward, L = Left, R = Right U = Up, D = Down) **I**) Representative putative GABAergic neuron aligned to the onset of licks after reward from mouse 2. **J**) Corresponding licks from the same behavioral session. **K-O)** Data from all 5 force sensors aligned to the onset of licks after reward (mouse 2). P) Correlation values

As was observed in the single-trial examples (Figures 3 & 4), the force signals that we measured coincided with licking behavior. To reveal the variability of these movements across many trials, we aligned the neural activity of a putative GABAergic neuron to reward and then sorted both the force measures and the neural data by the latency to the first lick following reward (Figures 5B & 5C). An examination of all the force sensor raster plots revealed high trial-to-trial variability when aligned to reward, even in well-trained mice. At the same time, the signals showed a consistent pattern as the mice anticipated and consumed the reward, likely reflecting the whole-body movements produced by the head-restrained mouse (forward, leftward, and downward movements; Figures 5D-H). The single unit activity was highly correlated with the movements (Figure 5P). We observed a similar pattern in a second neuron (Figure 5I) from a different mouse. In this example, the mouse moved forward, leftward, and upward, pushing down with both feet while consuming the reward (Figures 5 J-O). Interestingly, in this example, the load cells could register an oscillatory signal in the mouse’s movements during reward consumption (Figures 5J & 5K), which was most likely due to licking, which is well-known to have a stereotyped frequency at 5-8 Hz in mice. This pattern was also reflected in the activity of the isolated single unit (Figure 5I). This feature would have been hidden if only the average activity across trials is shown.

These preliminary results reveal the importance of recording the continuous movement of head-fixed animals. The neural activity more closely corresponds to the animals’ continuous movements, rather than to the events defined and imposed by the experimenter. Indeed, we found high correlations of the neural activity with load sensor signals (Figure 5P).

### Behavior during Pavlovian conditioning

Many studies in head-fixed mice employ Pavlovian conditioning approaches to study learning and to isolate specific goal-directed behaviors (Figure 6A, left). Virtually no study has examined the head movements or body forces exerted by head-fixed mice during such tasks. To demonstrate the utility of our device for such studies, we recorded VTA activity from two different well-trained mice performing a standard Pavlovian trace conditioning task (Figure 6A, right). During this task, mice received a 5 µl drop of sucrose (US) that was preceded by a 100-millisecond tone (CS) separated by a 1 second delay. Trials were separated by random inter-trial intervals (3-60 seconds). We simultaneously recorded neural activity and the forces exerted (Figures 6B-E). In this task, the mice displayed two phases of force exertion: a large movement immediately following reward and more subtle movements that occurred following the tone presentation (Figures 6B & 6D). Significantly, when we sorted the trials according to force initiation, the neural activity often closely followed the force changes, though each neuron that we recorded from showed distinct relationships to movements during each event (Figures 6B-E). Two neurons predominantly responded to movements following the tone (Figure 6C), and an additional two cells were sensitive to movements generated following the tone and reward (Figure 6E). Importantly, averaging the neural activity only conveyed the responses of neurons to movements following reward, but did not reveal the relationships that these neurons had with movements following the tone. Only by examining individual trials and sorting the data according to the continuous behavioral measures can these significant relationships be revealed clearly.

**Fig. 6.**
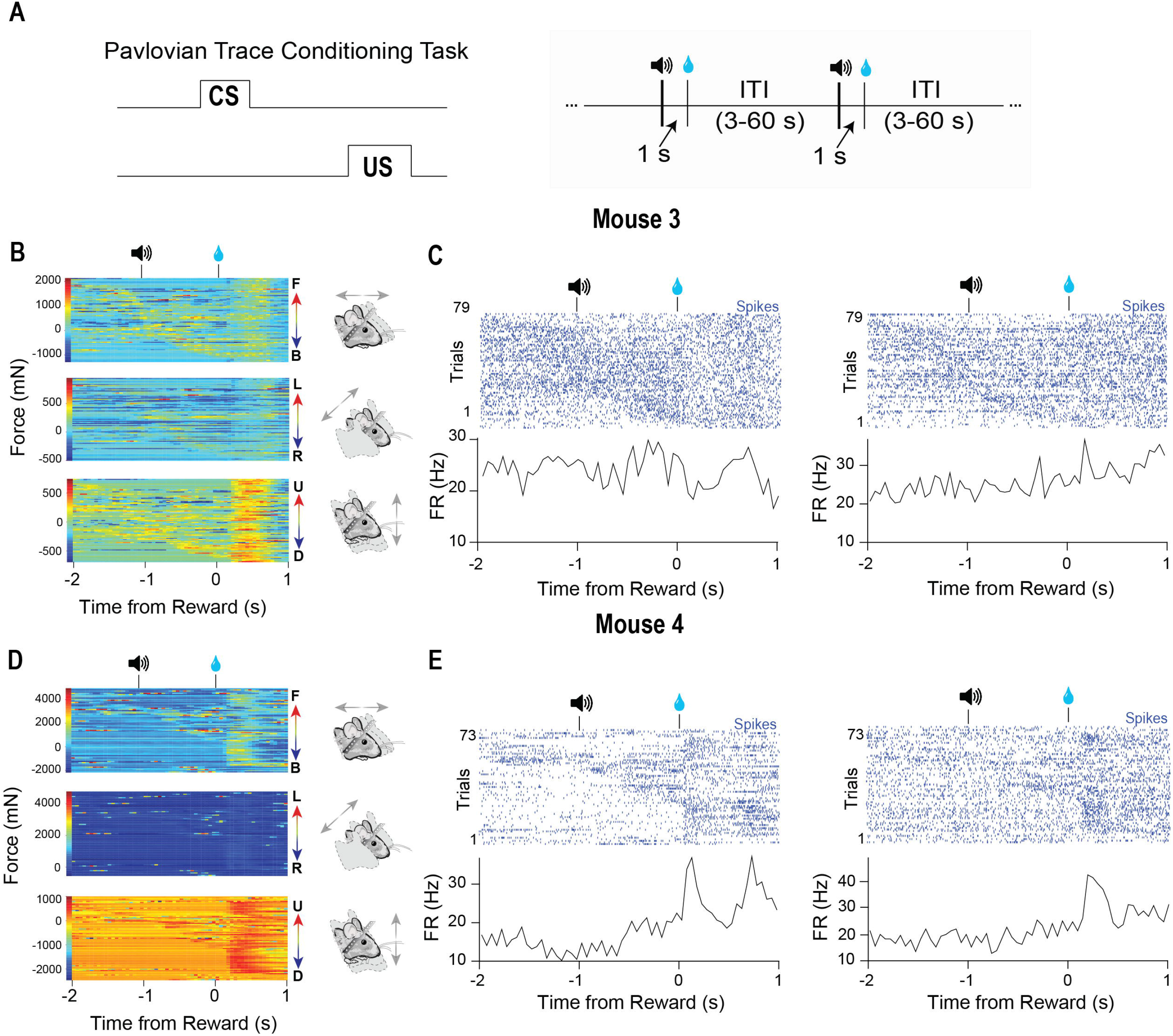
Characteristic force measures and neural activity on a Pavlovian conditioning task. **A)** Mice were trained on a Pavlovian trace conditioning task where a 100 ms tone (CS) was presented, followed by a 1 second delay and the delivery of a 10 % sucrose reward (US). There was a random interval trial interval (ITI) of 3-60 seconds. **B)** Head force measures showing mice exert large head forces during the task. **C)** Two corresponding representative putative GABAergic neurons from the VTA of mouse 3. Trials are sorted according to the start of the first forward movement. **D)** Head force measures during the task. **E)** Two corresponding representative putative GABAergic neurons from the VTA of mouse 4. Trials are sorted according to the start of the first forward movement.

## Discussion

One major limitation in behavioral neuroscience research is the lack of precise and continuous behavioral measurements. Head-fixation allows investigators to focus their analysis on a subset of movements, yet previous work using head-fixed animals largely ignored most of the actual behaviors exhibited. Our results contradict the common misconception that additional movements of the body and head are irrelevant once mice are head restrained. Ignoring these movements can limit our understanding of brain function. Our device reveals that even during head fixation, animals often make many continuous adjustments of the head and body while performing a reward-guided task. Furthermore, we show that neural activity can be closely related to these movements, revealing novel behavioral variables for explaining complex neural data.

Although head-fixation facilitates our ability to understand neural activity by providing precise measures of some behaviors such as licking or eye movements, our device shows that many head and body movements occur even though they are never explicitly measured (Figures 3 & 4). In addition, while clamped to a rigid structure, the strain on the head from trying to move the head presents a source of significant and highly variable sensory input. Therefore, two major confounds exist in previous studies using head-fixed approaches while investigating the neural substrates of a given behavior. If these considerations are not taken into account, one is prone to explaining neural activity with arbitrary, irrelevant, and unnecessarily complex variables.

We show that mice make many head and body movements during the performance of traditional head-fixation tasks, and their neural activity can follow these dynamics quite closely (Figures 3–6). Thus, many studies claiming to investigate the neural correlates of variables such as reward or value during head-fixed studies have failed to notice a large movement confound. It would be helpful to reexamine all previous head-fixed studies using continuous force measures introduced here.

Traditional *in vivo* electrophysiology in head-fixed animals usually relies on experimenter-defined behavioral events, rather than continuous processes (Yin, 2016;Yin, 2017). Actions are often assumed to be ‘all-or-none’ discrete events, such as reaching, licking, or saccades(Georgopoulos et al., 1986;Schultz et al., 1992;Hikosaka et al., 2000). Behavior is thus only quantified as a categorical time stamp, rather than as a continuous process which unfolds over time. As a result, investigators have only attempted to understand neural activity by aligning it to this observer-defined action and analyzing its activity around this time point. The neural activity that occurs in between experimenter-defined events is usually neglected. However, recent work has shown that continuous behavioral measurements with high spatial and temporal resolution can reveal previously neglected relationships between neural activity and behavior (Barter et al., 2014;Kim et al., 2014;Barter et al., 2015a;Barter et al., 2015b;Bartholomew et al., 2016;Hughes et al., 2019;Kim et al., 2019). Our system demonstrates that, with continuous behavior measures, neural activity outside of these observed defined time-points becomes intelligible.

Variability in neural activity has long been a puzzle for neuroscience (Shadlen and Newsome, 1998;Osborne et al., 2005;Churchland et al., 2010). The results obtained from our novel head-fixation device point towards a potential source of this variability: continuous behavior that has not been noticed or measured. By aligning the force measures and neural activity to experimenter-defined events, but sorting the trials according to continuous behavioral measures, the neural activity is shown to reflect the continuous force measures and not the events (Figures 5 & 6). Our results suggest that it is not noise but meaningful neural activity that must be explained by any theory of the brain.

While newer head-fixed set ups use a running ball or an accelerometer in order to measure the behavior, there are several disadvantages associated with these setups (Harvey et al., 2009;Coddington and Dudman, 2018). First, it is impossible to detect direction-specific adjustments of the head and body with either an accelerometer basket or a running ball. Second, movement of the ball is only an indirect measure of the actual kinematics. The ball has its own momentum that can be independent of the animal’s movement. For example, if the ball is moving to the left, but the mouse wants to go forward, many opposing adjustments of the limbs and body have to occur to counteract the ball’s momentum in order to provide forward motion.

The measures from the ball could indicate a turn from left to forward whereas the mice are making large adjustments in different directions in order to obtain the new heading. Our head-fixation system can be used in conjunction with treadmills balls to detect these adjustments, where the direction-specific head movements can be measured and directly compared with the coordinates obtained from the ball. This would make it possible to dissociate head movements and limb movements.

In summary, our head-fixation system can overcome many of the limitations in conventional head-fixed setups. The load cells allow direct and continuous measurement of the force exerted by the mouse’s head as well as the body. The chief advantages of our design are its low-cost (~$140, see supplementary materials for parts list), ease of construction, and high temporal and spatial resolution.

## Methods

### Mice

All experimental procedures were approved by the Mouse Care and Use Committee at Duke University. Four C57BL/6J mice between 2-6 months old were used for experiments (Jackson Labs, Bar Harbor, ME). Mice were housed in groups of 3-4 mice per cage on a 12:12 light cycle, with experiments conducted during the light phase. During days where mice were run on experiments, they were water restricted and maintained at 85-90% of their initial body weights. Mice received free access to water for approximately 2 hours following the daily experimental session.

### Load Cells and Circuit

Each load cell (RobotShop, Swanton, VT, USA) is a full Wheatstone bridge with a configuration of four balanced resistors based on a known excitation voltage, as shown below:

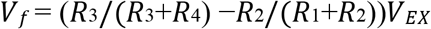

Where R is the resistor, *V*_*EX*_ is a known constant excitation voltage, and output *V*_*f*_ is variable depending on the shape of the strain gauge. Thus, a bidirectional sensing amplifier connected to the load cell will output 2.5V at zero load. When the load is increased or decreased, the voltage will correspondingly increase or decrease. This relationship is described by the following:

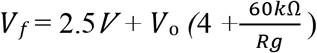

Where *Rg* is the resistance of the potentiometer, and Vo is the voltage output. The sensitivity of the circuit can be adjusted using the potentiometer, which is a precise trimmer potentiometer with 200 Ohms resistance. The amplifier gain (*Gf*) from this chip is described as below.

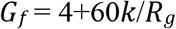

The voltage from the amplifier (INA125UA, Texas Instruments) is then linearly correlated with the force on the load cell. That means we can use the voltage to describe the force exerted on each orthogonal load cell. From Newton’s second law (*F = ma*, where *F* = force, *m* = mass and *a* = acceleration), a mass with a known quantity placed on the load cell can be multiplied by the gravitational constant to obtain a force-voltage conversion factor (F/V). This conversion factor can then be used to convert the voltage to force exerted. The resistance values for load cells used for behavioral experiments were: 103 Ohms (F/B force sensor); 102.5 Ohms (S/S Force sensor); 103 Ohms (U/D force sensor); 102.8 Ohms (right feet force sensor); 103.1 Ohms (left feet force sensor).

### Head-fixed behavioral setup

The apparatus was 3D printed in PLA plastic using a MakerGear M2 3D printer (MakerGear, Beachwood, OH, USA). Both mouse perch and head fixation apparatus were bolted to a steel base plate (Thorlabs, Newton, NJ, USA), and elevated to give sufficient space for the reward delivery apparatus. In the floor of the perch, two 100g load cells (RobotShop, Swanton, VT, USA) were used to measure ground reaction forces on the left and right side of the mice. The mice’s head implants were clamped to a custom frame that was suspended above them. In the head fixation apparatus between the point of head fixation and the base plate, three 100g load cells of the same type were arranged orthogonally to detect force changes in three dimensions (forwards-backwards, up-down, and left-right). These load cells were connected in series and arranged close together and in line with the mouse’s head to minimize twisting forces. Each load cell was connected by a shielded cable to an amplifier circuit, which was in turn connected to the data acquisition system. Each load cell signal was recorded continuously as an analog voltage. Load cells translate mechanical stress caused by an applied load into a voltage signal. Proper grounding of the circuit is important, as it can introduce noise into electrophysiological and force sensor recordings. A 10 % sucrose solution gravity-fed from a reservoir was delivered to the mice via a metal spout placed in close proximity to their mouth. Sucrose delivery was controlled through the use of a solenoid valve (161T010, NResearch, NJ). A capacitance-touch sensor (MPR121, AdaFruit.com) was clamped to the metal spout in order to record licking behavior. All analog voltage signals (1 Khz) and electrophysiological data were recorded using a Blackrock Cerebrus recording system (Blackrock Microsystems) for offline analysis. Electrical noise that appeared in force sensor signals was removed in post-processing in some sessions.

### *Wireless* in Vivo *Electrophysiology*

Mice were anesthetized with 2.0 to 3.0% isoflurane (David Kopf Instruments, Tujunga, CA), maintained at 1.0 to 1.5 % isoflurane for surgery using a stereotactic frame, as previously described(Fan et al., 2011;Fan et al., 2012). A craniotomy was then made above the VTA, and fixed 16-channel electrode arrays, with tungsten electrodes in a 4 x 4 configuration (35 μm diameter, 150 μm spacing, 5 mm length, Innovative Neurophysiology, Inc.) were lowered into the VTA (AP: 3.2 mm, ML 0.5 mm, DV, −4.2 mm). After electrodes were inserted, they were secured to the skull using screws and dental acrylic, followed by the insertion of a metal bar around the dental acrylic to allow for head-fixation (Toda et al., 2017).

### Behavioral tasks

Mice were allowed to recover for 2 weeks, then mice (*n* = 2) were trained on a fixed time (FT) reinforcement schedule for approximately 1-2 weeks (Rossi et al., 2013;Toda et al., 2017). An additional 2 mice were trained on a trace Pavlovian conditioning task for approximately 5-7 days until anticipatory licking was observed after the sound of the tone. The tone lasted 100 ms, followed by a 1 second delay. Reward was then delivered after the delay. A random ITI of 3-60 seconds was used in between reward trials. A miniaturized wireless head stage (Triangle Biosystems) that communicated with a Cerebrus data acquisition system (Blackrock Microsystems) was used to record electrophysiological data (Fan et al., 2011;Barter et al., 2014). All single unit data were then sorted offline using OfflineSorter (Plexon). All plots for electrophysiological and force signal data were created with NeuroExplorer (Nex Technologies) using 50 ms bins. In order to be included for analysis, neural data must have had a 3:1 signal-to-noise ratio, with an 800 μs or greater refractory period.

## Supporting information

construction video back

Construction video front

## Acknowledgments

This study was supported by NIH grants NS094754, DA040701, and MH112883 to HHY. We would like to dedicate this paper to the memory of Kobe Bryant (1978-2020), whose artistry and dedication have been a major source of inspiration.

