## Supplemental information for "A head-fixation system for continuous monitoring of force generated during behavior"

*Assembly Instructions:*

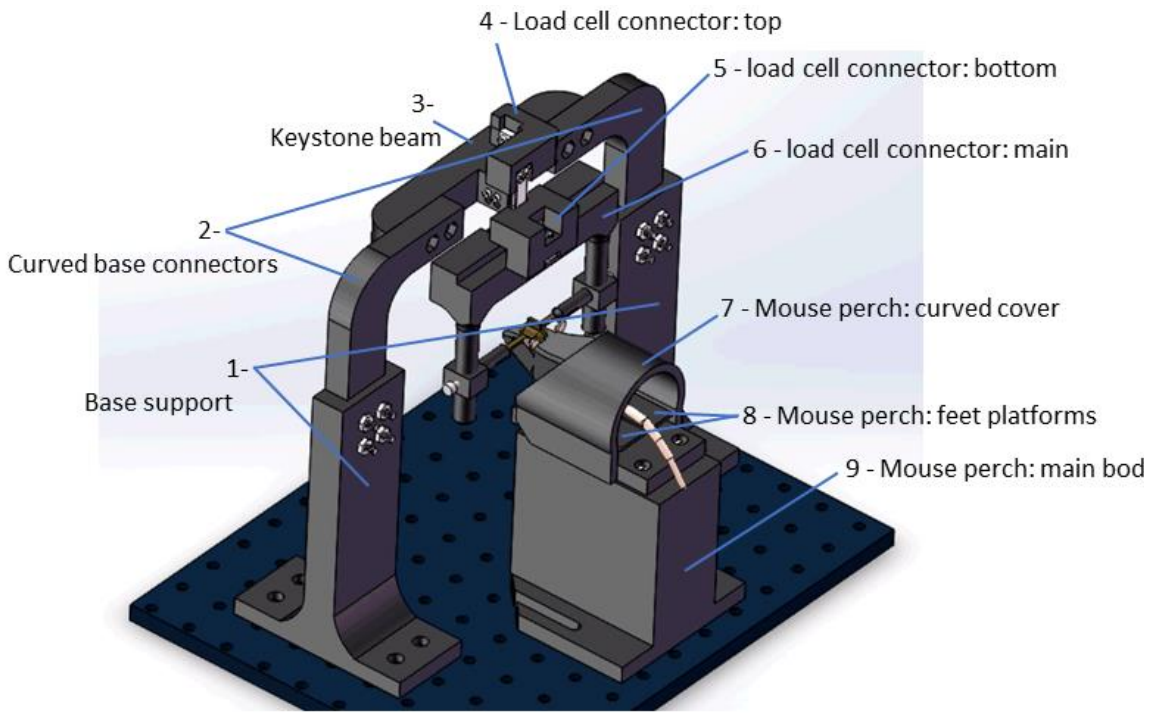


Instructions for assembling the head fixation apparatus:

1) Attach base supports (parts 1) to the steel baseplate using M6 bolts.

2) Attach curved base connectors (parts 2) to base support (parts 1) using M6 nuts and bolts.

3) Attach keystone beam (part 3) to base connectors (parts 2) using M6 nuts and bolts.

4) Attach the lower end of a load cell to keystone beam (part 3) using M2.5 bolts.

5) Attach the upper end of the load cell to the thin side of top load cell connector (part 4) using M2.5 bolts.

6) Attach the upper end of a second load cell to the thick side of the top load cell connector (part 4) using M2.5 bolts.

7) Attach bottom load cell connector (part 5) to the lower end of the second load cell using M2.5 bolts.

8) Attach a third load cell to main load cell connector (part 6) using M2.5 bolts.

9) Attach the other side of the third load cell to bottom load cell connector (part 5) using M2.5 bolts.

Instructions for assembling the perch apparatus:

1) Attach fourth and fifth load cells side by side to the inside of the curved cover of the mouse perch (part 7) using M2.5 bolts.

2) For both the fourth and fifth load cell, attach the free end to a copy of the platforms of the mouse perch (parts 8) using M2.5 bolts. The flat side of feet platforms of the mouse perch (parts 8) should face upward.

3) Attach the curved cover of the mouse perch (part 7) to the main body of the mouse perch (part 9) using M6 bolts. Attachment holes in part 9 should be self-threading.

4) Attach the main body of the mouse perch (part 9) to the steel baseplate using M6 bolts through the attachment channels at the base of the main body of the mouse perch (part 9).

*See supplemental video S1 and S2 for a 3D rendering of construction.

*Parts list*:

1) Spool of 3D printer filament. Any rigid plastic works. We used PLA on Fused deposition modeling (FDM) printer (1x $18.00 per spool, https://www.amazon.com/SUNLU-Filament-1-

75mm-Printer-Black/dp/B07Q82HVTT/ref=sr_1_14?keywords=PLA+filament&qid=1579894242&s=industrial&sr=1-14).

2) 100g load cell (5x, $7.00 each, https://www.robotshop.com/en/100g-micro-load-cell.html?gclid=EAIaIQobChMIybPxse2c5wIVzJ-zCh353gR0EAQYBSABEgJbLvD_BwE).

3) INA125 Instrumentation Amplifier (5x, ~$7.00 each, https://www.digikey.com/product-detail/en/texas-instruments/INA125UA/INA125UA-ND/300986).

4) Trimmer potentiometer (5x, $1.36 each, https://www.digikey.com/product-detail/en/nidec-copal-electronics/ST4ETB101/ST4ETB101CT-ND/738511).

5) 4 pin male connector (5x, $0.4 each, https://www.digikey.com/product-detail/en/1718560004/WM10155-ND/4423111/?itemSeq=314595070).

6) 4 pin female connector (5x, $0.45 each, https://www.digikey.com/product-detail/en/PPTC041LFBN-RC/S7002-ND/810144/?itemSeq=314595184).

7) 2 pin male connector (6x, $0.2 each, https://www.digikey.com/product-detail/en/0022280020/WM19475-ND/3157814/?itemSeq=314595790).

8) Jumper wires assortment (1x, $7.29 each,https://www.amazon.com/Yueton-Multicolored-Female-Breadboard-Jumper/dp/B01DDD1LXU/ref=sr_1_25?keywords=jumper+wires&qid=1579893448&s=hi&sr=1-25)

9) 3-conductor shielded cable for connecting load cells to amplifier circuits (~$60 for 3.05m, https://www.digikey.com/product-detail/en/alpha-wire/85243CY-BK005/85243CYBK005-10-ND/9090384 )

10) M6 nuts and bolts assortment (1x, $19.50, https://www.amazon.com/120Pcs-Stainless-Socket-Washers-Assortment/dp/B01N0ZU46G?ref_=fsclp_pl_dp_15)

11) M2.5 bolts assortment (1x, $9.99, https://www.amazon.com/HVAZI-Stainless-Phillips-Machine-Assortment/dp/B075QKZ8PY/ref=sr_1_7?keywords=m2.5+bolts&qid=1579890750&s=hi&sr=1-7)

12) Printed circuit board (1x, ~$3.00 per board, depending on supplier and quantity).
